# Cerebellar influences on neocortical development in humans and mice

**DOI:** 10.64898/2026.08.19.745702

**Authors:** Carolin Gaiser, Nathaniel Germain, Thomas Jacobs, Maarten A. Frens, Jörn Diedrichsen, Jeremy Labrecque, M. Mallar Chakravarty, Gabriel A. Devenyi, Aleksandra Badura, Ryan L. Muetzel

## Abstract

The cerebellum has long been considered a late-maturing structure subordinate to neocortical development, therefore its potential role as an early driver of cortical organization remains largely unexplored. Using two large longitudinal neuroimaging cohorts of developing children together with lesion experiments in mice, we show that early cerebellar morphology may drive neocortical maturation in a regionally specific manner. These cross-species findings implicate the cerebellum as a possible regulator of neocortical organization.

## Main Text

An emerging but underappreciated body of evidence hints at a broader organizational role for the cerebellum in shaping brain structure and function. Despite pronounced differences in size, the cerebellar and cerebral cortices scale in a tightly coordinated manner across mammalian species, maintaining a near-constant neuronal ratio of ∼3.6 cerebellar to 1 neocortical neuron^1^. This conserved scaling relationship suggests concerted evolution of the two structures.

During human development, the cerebellum and cerebral cortex likewise exhibit parallel maturation patterns, with motor regions developing earlier and higher-order cognitive regions maturing into late adolescence^2,3^. Notably, cerebellar development appears to slightly precede cortical maturation^4^, suggesting a potential upstream role of the cerebellum during neurodevelopment. Consistent with this idea, clinical observations in preterm infants demonstrate that impaired cerebellar growth during the third trimester is associated with reduced growth of uninjured, contralateral cortical regions that are targets of cerebellar projection pathways later in development^5^.

This potential influence of cerebellar growth on cortical development is supported by known extensive cerebrocerebellar connectivity. Cerebellar output influences cortical activity through well-defined thalamic pathways, and experimental animal studies demonstrate that cerebellar signaling can modulate cortical plasticity in both sensory and associative regions^6^. Moreover, cerebellar-prefrontal circuits have been shown to regulate higher-order behaviors, with targeted cerebellar manipulations inducing, and rescuing, social and repetitive behavioral deficits by modulating altering cortical function even beyond early developmental periods in an *Autism Spectrum Disorder* (ASD) mouse model^7^. Together, these findings suggest that the cerebellum might guide the maturation of remote neural circuitry and influence cortical development.

Despite this converging evidence from evolution, clinical reports and animal work, it remains unclear whether differences in cerebellar maturation influence subsequent cortical developmental trajectories. In the present study, we therefore combine large-scale neuroimaging datasets of developing children and adolescents with mouse lesion models to test the causal role of the cerebellum in neocortical maturation.

Using longitudinal structural *magnetic resonance imaging* (MRI) data from the two largest prospective population-based cohorts of brain development: the Adolescent Brain Cognitive Development (ABCD) study^8^ and the Generation R study^9^, we tested whether cerebellar morphology in childhood drives subsequent cortical maturation. Participants with repeated neuroimaging at baseline and follow-up were included (Fig. 1a).

**Figure 1.**
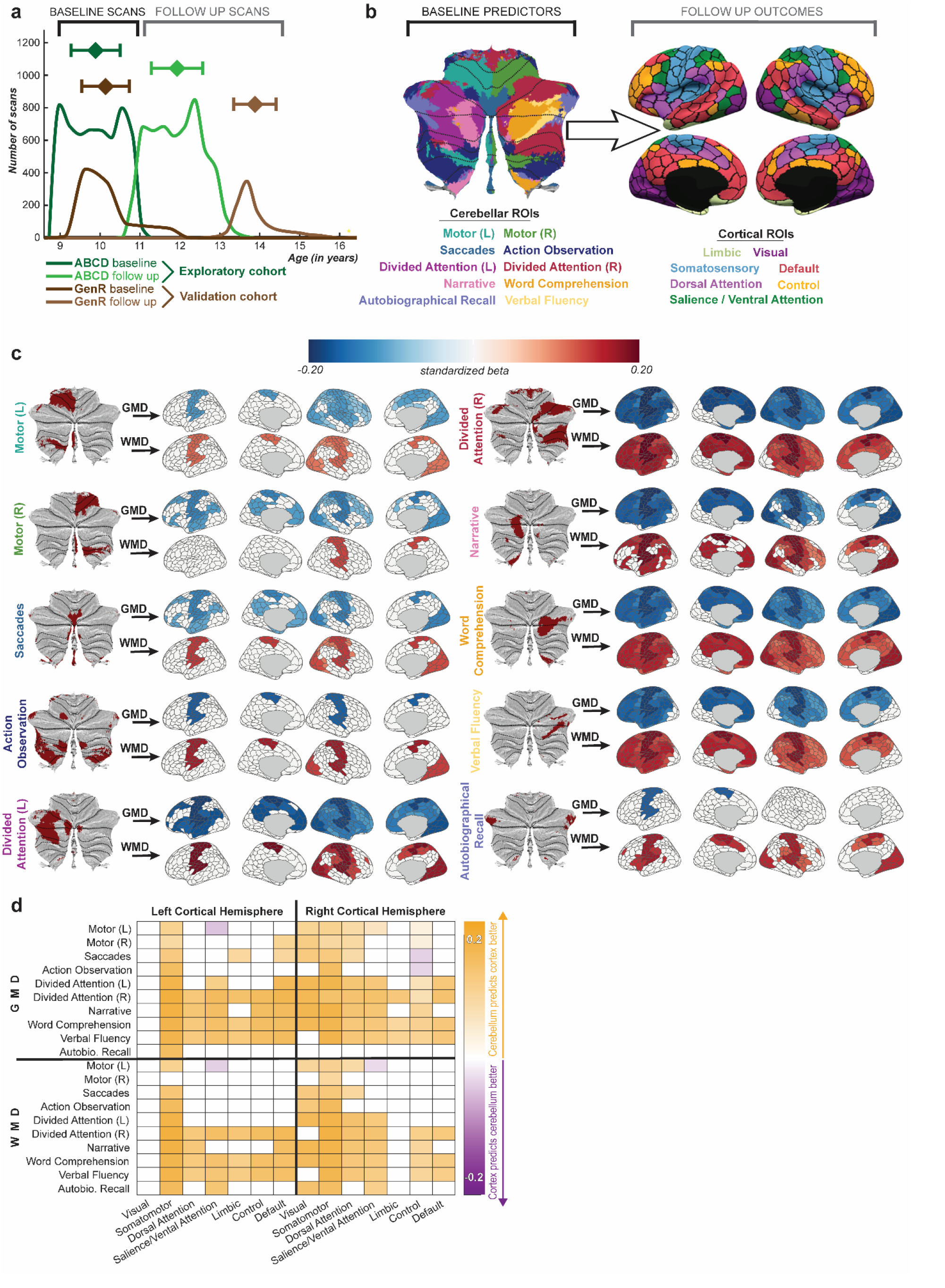
Cerebellar morphology in childhood drives subsequent cortical maturation. **a**. Overview of the prospective, population-based human imaging datasets used. In shades of green, the histograms of the ABCD study participants (N = 5,310) included in this study at baseline (mean age at scan = 9.9 years, range = 8.9 - 11.0) and follow up (mean age at scan = 11.9 years, range = 10.6 - 13.8). In shades of brown, the histograms of the Generation R study participants (N = 968) included in this study at baseline (mean age at scan = 10.1 years, range = 8.9 – 12.0) and follow up (mean age at scan = 13.9 years, range = 12.6 - 16.7). The larger ABCD cohort was treated as the exploratory sample and the Generation R cohort as a validation sample. **b**. Predictor and outcome regions-of-interest (ROIs) based on functionally defined atlases. Cerebellar functional ROIs are based on the Multi-Domain Task Battery^11^ and cerebral functional ROIs are based on the Yeo 7-network parcellation^10^. **c**. Results showing the effect of baseline *Grey Matter Density* (GMD) and *White Matter Density* (WMD) in each of the cerebellar functional ROIs on changes in cerebral thickness. Blue indicates a significant decrease and red a significant increase in cortical thickness. **d**. Directionality of cerebellar–cortical associations. Matrix displays the net standardized regression coefficient (β_cerebellum→cortex - β_cortex→cerebellum) for all significant cerebellar-cortical pairs. Yellow indicates a stronger association between baseline cerebellar morphology and subsequent cortical maturation; purple indicates a stronger association between baseline cortical morphology and subsequent cerebellar maturation. Only associations surviving FDR correction in both cohorts independently are shown.

We modeled the annualized rate of change in cortical thickness across seven cortical network regions^10^ as a function of baseline cerebellar *grey matter density* (GMD) and *white matter density* (WMD) in ten functional cerebellar *regions-of-interest* (ROIs)^11^ (Fig. 1b). Models were adjusted for biological sex, age at baseline, gestational age at birth, a comprehensive set of behavioral, health, maternal, and socioeconomic factors (see Supplementary Information for complete list), as well as total brain volume at baseline. Further, for each cortical region, the baseline regional measure was included in the model, allowing us to interpret the model as a change in cortical structure over time that cannot be accounted for by the cortex’s own baseline morphology. To ensure robustness and generalizability, we adopted a two-stage approach: the larger ABCD cohort (N = 5,310) served as an exploratory sample, and Generation R (N = 968) as an independent validation cohort. Associations were considered significant only if observed in both cohorts independently after false discovery rate (FDR-BH) correction. We additionally applied the same models to cerebellar^11^, cortical^10^, and subcortical^12^ volumes, however fewer significant associations were observed after total brain volume correction (Supplementary Fig. 1A and 1B).

Results show that cerebellar morphology was robustly associated with cortical maturation in a regionally specific manner. Higher cerebellar GMD at baseline was generally associated with decreases in cortical thickness at follow up, whereas higher cerebellar WMD was associated with increases. Notably, this relationship was topographically specific: cerebellar motor regions were associated mostly with maturation of sensorimotor cortical areas, while posterior cognitive cerebellar regions were associated with widespread cortical regions (Fig. 1c).

To assess the directionality of these associations, we applied the same models in reverse, using baseline cortical thickness to examine subsequent regional cerebellar changes. We then compared standardized betas from the initial models (cerebellum→cortex) and reverse models (cortex → cerebellum) by subtracting the latter from the former. Across nearly all significant associations, the cerebellar-to-cortical association was consistently stronger than the reverse, providing temporal evidence that these relationships are predominantly driven from cerebellum to cortex (Fig. 1d).

Moving from observational human data to experimental data, we next used a clinically relevant rodent model of early cerebellar damage to directly test causality. Guided by the directional associations in our human cohorts, we induced focal lesions of the right Crus I at postnatal day (P)14 or P21, periods of heightened developmental vulnerability. Outcomes were compared to age-matched sham-operated controls, alongside a non-surgical group to account for the effects of anesthesia and surgical stress.

We used a *deformation-based morphometry* (DBM) pipeline to assess voxel-wise morphological differences across subjects^13^. This approach employs an unbiased group average rather than a predefined template. Individual deviations from this average were quantified and compared at the group level. Statistical analysis using linear models with group and sex as covariates, with false discovery rate correction for multiple comparisons, revealed significant regional differences in all lesion groups. In contrast, sham groups showed no significant or spatially coherent effects (Supplementary Table 1; Supplementary Fig. 2). The non-surgical group served as the reference and is therefore not shown.

Unilateral right Crus I lesions induced widespread structural alterations extending beyond the cerebellum into sensorimotor, visual, cingulate, and somatosensory cortical regions, as well as connected subcortical structures (Fig. 2b–d). Although both lesion groups showed overlapping deformation patterns (Fig. 2d), earlier lesions at P14 produced broader, larger, and more bilateral effects than lesions at P21, as seen in expansions found in the visual and somatosensory cortices. This pattern is consistent with greater sensitivity of earlier developmental stages and a potential sensitive-period effect in cerebello-cortical development^15^. Several regions that showed no or only subtle alterations following the P21 lesion, were selectively altered only in the P14 group, such as the expansions in the contralateral temporal association cortex, as well as deformations in non-cortical structures (Fig. 2 c,d; Supplementary Figure 2).

**Figure 2.**
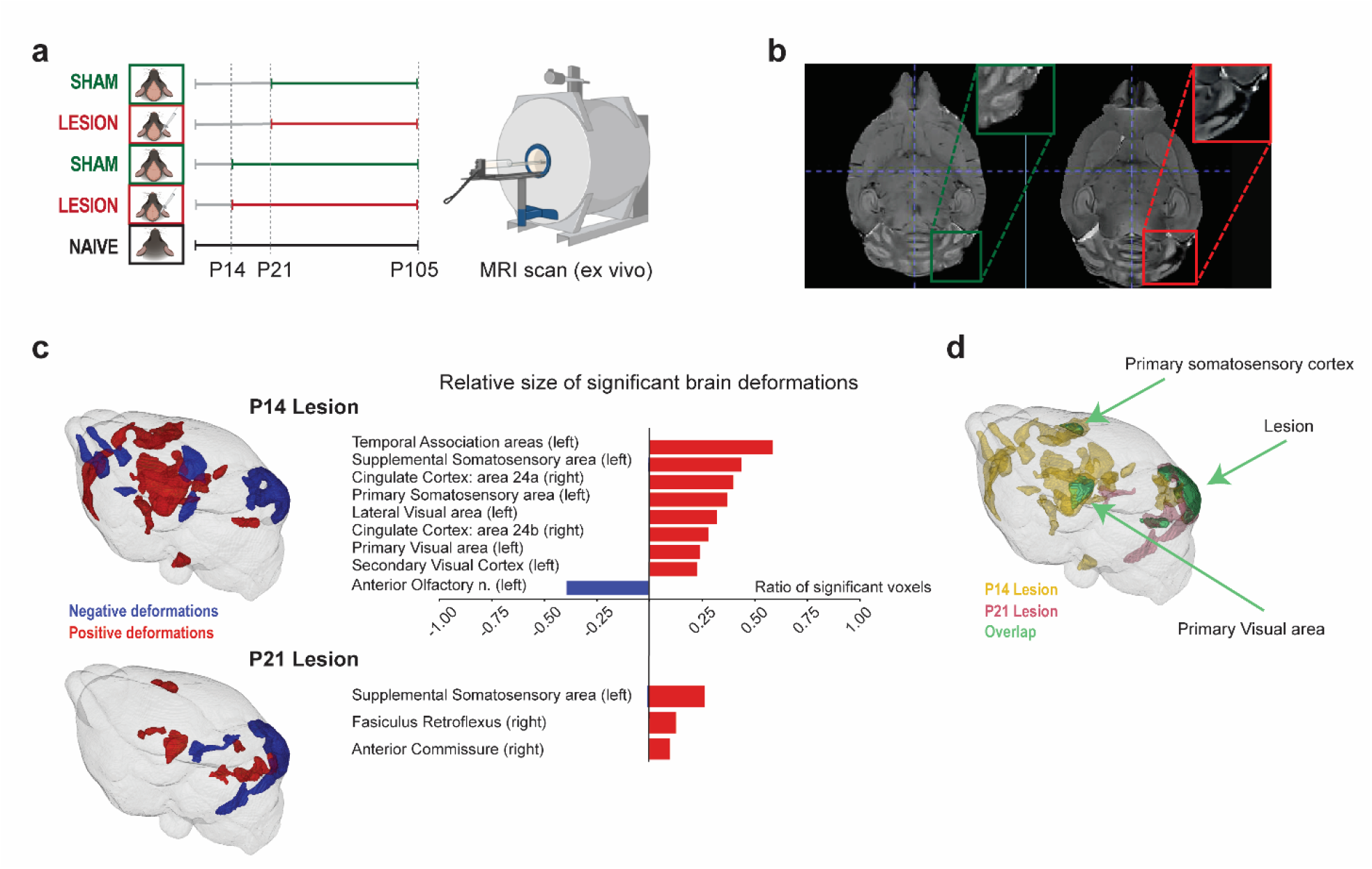
Early postnatal Crus I lesions in mice induce age-dependent brain-wide structural alterations. **a**. Experimental timeline and lesion paradigm targeting Crus I at either P14 or P2. **b**. Axial MRI slices of a sham control (left) and lesioned mouse (right); insert shows the extent of the lesion spanning Crus I. **c**. Brain-wide deformation maps, represented in glass brain renders, as well Allen Brain Atlas^14^ structure level analysis of significant voxel localization, revealed distributed structural alterations following Crus I lesions, including both bilateral and contralateral changes across cortical and subcortical regions beyond the cerebellum. Lesions at P14 produced broader and more pronounced effects than lesions at P21, particularly within neocortical areas. Red indicates relative volume increases, and blue decreases (5–1% FDR corrected) **d**. Glass brain renders of significant voxels in both the P14 (in yellow) and P21 (in red) lesion group, as well as their overlap (in green), clearly shows the lesion and revealed that the visual and somatosensory areas are affected in both groups. By contrast, many of the profound changes observed in the P14 lesion group such as expansions in the ipsilateral cingulate cortex or the contralateral temporal association cortex do not occur when the lesion takes place at P21.

Together, these results across human and rodent data demonstrate the causal influence of the cerebellum in shaping neocortical development. Importantly, across species, cerebellar effects were aligned with known structural and functional cerebrocerebellar circuits. In humans, sensorimotor cortical regions were influenced by cerebellar indicators across almost all functional subregions, whereas posterior/cognitive cerebellar regions showed more widespread associations across the neocortex, in line with known functional connectivity^16^. In mice, focal lesions of right Crus I preferentially affected the contralateral somatosensory cortex and visual areas, with additional effects in temporal association and cingulate regions when the lesion occurred earlier (P14), consistent with established cerebellar projection pathways^17, 18^. Earlier lesions produced larger cortical effects, highlighting a developmental window during which cerebellar integrity is particularly important.

These findings fit well with theoretical accounts proposing that the cerebellum acts as a coordinator of task-specific neuronal activity, integrating inputs from distributed cortical systems according to a shared computational principle rather than domain-specific content^19, 20^. From this perspective, cerebellar regions engaged in sensorimotor versus cognitive processing would be expected to differentially influence the maturation of their connected cortical targets, as observed here across species and developmental stages. Disruption of cerebellar development may therefore propagate through cerebrocerebellar circuits to alter cortical maturation remotely, consistent with the concept of cerebellar diaschisis^15^.

These findings also provide a developmental context for clinical observations that posterior cerebellar damage preferentially affects executive, affective, and social functions, as described in the cerebellar cognitive affective syndrome^21^. More broadly, they extend prior work by showing that cerebellar contributions to cortical organization are not limited to mature brain function but emerge during development, with cerebellar maturation preceding and predicting cortical change more strongly than the reverse. This is consistent with evolutionary evidence showing disproportionate expansion of the cerebellum in apes and humans, where cerebellar specialization may have supported the emergence of complex cognitive behaviors^22^.

Taken together, these findings position the cerebellum as a potential driver of cortical developmental trajectories in a regionally and temporally specific manner and highlight the need to consider cerebrocerebellar interactions as a fundamental organizing principle of brain development across species.

## Online Methods

### Data

#### Human Cohorts

Neuroimaging scans of the two largest prospective cohorts of child development are used, the *Adolescent Brain Cognitive Development* (ABCD) study^8^ and the Generation R study^9^.

##### ABCD

The ABCD Study examines brain development from pre-adolescence into adulthood. This large-scale, ongoing project is conducted across 21 data acquisition sites in the United States and recruited approximately 11,500 children at baseline, who are followed for ten years with extensive assessments at multiple timepoints. The baseline cohort, recruited between September 1, 2016, and August 31, 2018, consists of nine- and ten-year-old children and their parents or guardians. Participants complete annual laboratory-based assessments, including MRI every two years. The brain imaging data were obtained from the baseline wave and two-year follow-up assessments (release 5.0). Recruitment primarily occurred through public elementary schools (including charter schools) and private schools. Ethical approval was received from the institutional review boards of the University of California (San Diego) and each ABCD site, adhering to their Institutional Review Board approved protocols, state regulations, and local resources. Informed consent has been received from the included participants.

##### Generation R

The Generation R Study is a population-based prospective cohort study that follows participants from fetal life onward. Between 2002 and 2006, 9,778 pregnant women living in Rotterdam, The Netherlands, were enrolled. Extensive data have been collected from both children and their caregivers at multiple timepoints, including MRI. For the present study, MRI data from the second measurement wave (mean age = 10.2 years) and the third measurement wave (mean age = 14 years) were used.

Written informed consent from both parents and assent from all participants was obtained, and the study was approved by the Medical Ethical Committee of the Erasmus Medical Center. Participants did not receive monetary compensation, but their travel costs were reimbursed. Additionally, as a token of appreciation for their participation, they received small gifts valued at 10€ or less, such as a drinking bottle, a bag, a power bank, or similar items.

#### Mice

Ethical approval was granted prior to the experiments (by Centrale Commissie Dierproeven (CDC), the Hague, the Netherlands), as required by Dutch law. All experiments were performed according to institutional, national and European Union guidelines and legislation, AVD10100202115462.

During this study both female and male C57BL/6J (later referred to as Black 6) mice were used. The mice were socially housed in environmentally enriched cages (T-3 IVC). Each cage contained 2-3 mice and included a running wheel, a cardboard house, and additional nesting material. Mice were housed together after weaning at P21 with mixed sham and lesion groups. Mice were housed under a 12-hour light/dark cycle and provided with ad libitum food and water. Temperature and humidity were controlled.

### Mice lesions and protocol

This study uses a somatic lesion model, in which we lesion the right cerebellar hemisphere, specifically lobule Crus I at P14 and P21 using the same procedure. All mice were fixated on a stereotaxic frame (Kopf Instruments), on a temperature-controlled bed. We applied general anesthesia consisting of 1-5% isoflurane (ISOFLUTEK 1000) together with subcutaneous injection of Buprenorphine (Bupredine,AST Farma) and Carprofen (Caraporal, AST Farma). After fixation and induction of the anesthesia, an incision was made to expose the occipital muscle and bone. Lidocaine (AST Farma) drops where applied on both the occipital muscle and bone for local sedation. Muscles directly above Crus I were removed and a narrow craniotomy was made using a sharp forceps. In the mice that were part of the lesion group, the right Crus I region was removed using aspiration. To stop bleeding, a hemostatic sponge (Spongostan, Ethicon) was placed on the brain, covering the exposed brain. Sham conditioned mice did not undergo lesioning of the right Crus I region but did undergo all other surgical procedures. The surgery was completed with the closing of the skull with soluble sutures (vicryl 6-0, Ethicon) and tissue adhesives (Vetbond, 3M).

## Neuroimaging acquisition and preprocessing

### Human Cohorts

#### Image Acquisition

High resolution T^1^-weighted MRI scans were acquired on multiple 3T scanners in the ABCD study (Siemens Prisma, *General Electric* (GE) 750, and Philips), and on a study-dedicated 3T scanner (GE MR750w) in the Generation R study using the following parameters (listed per scanner): Siemens Prisma (ABCD): T^R^ = 2500 ms, T^E^ = 2.88 ms, T^I^ = 1060 ms, flip angle = 8°, field of view = 256 ×256 mm; GE 750 (ABCD): T^R^ = 2500 ms, T^E^ = 2.00 ms, T^I^ = 1060 ms, flip angle = 8°, field of view = 256 ×256 mm; Philips (ABCD): T^R^ = 6.31 ms, T^E^ = 2.90 ms, T^I^ = 1060 ms, flip angle = 8°, field of view = 256 ×240 mm; GE MR750w (Generation R): T^R^ = 8.77 ms, T^E^ = 3.4 ms, T^I^ = 600 ms, flip angle = 10°, acquisition time = 5 min 20 s, field of view = 220 ×220 mm.

All scans were acquired in 1x1x1 mm^3^ isotropic resolution. The full details on image acquisition in ABCD and Generation R were described in previous studies^23, 24^.

#### Image Preprocessing

The identical preprocessing pipeline, SMRIPrep^25^, on the same high-performance computing system was used for both the ABCD and Generation R data. Briefly, SMRIPrep encompasses surface reconstruction using FreeSurfer (version 6.0.0) and non-brain tissue was removed, voxel intensities were adjusted for B^1^ inhomogeneities, and images were linearly and nonlinearly registered to standard stereotactic space (MNI152 NLin2009cAsym 1x1x1 mm resolution) using ANTs (github: https://github.com/ANTsX/ANTs.git). The tissue segmentation procedure (FSL FAST: https://fsl.fmrib.ox.ac.uk/fsl/fslwiki/FAST) resulted in binary classifications of voxels, as well as in per-voxel tissue class probability estimates. These probability estimates can be interpreted as the likelihood of a given voxel being grey matter, white matter, or cerebrospinal fluid. Further, the nonlinear registration produced a nonlinear warp file (which included the linear initialization) from which we calculated the determinant of the Jacobian matrix for each voxel. This determinant was used as a measure of volume of that voxel relative to its volume in standard stereotactic space for calculations of volumes in the cerebellar ROIs.

#### Regions-of-Interest (ROIs)

We parcellated functional ROIs of the cerebellum using the Multi-Domain Task Battery atlas^11^ as well as of the neocortex using the 7 network Schaefer atlas^10^. For cerebellar ROIs, scans were normalized to the MNI-aligned atlas template and mean *grey matter density* (GMD), *white matter density* (WMD), and volumes (defined as sum of Jacobian determinants) were extracted for each of the ten functional regions. For neocortical ROIs, the Schaefer/Yeo 400-parcel, 7-network parcellation was applied to individual cortical surfaces registered to fsaverage space by sMRIPrep. Mean cortical thickness, surface area, and volume were extracted per parcel by averaging across vertices using FreeSurfer. Furthermore, hemispheric volumes of FreeSurfer-based subcortical structures^12^ (Thalamus, Caudate, Putamen, Pallidum, Hippocampus, Amygdala, Accumbens, Ventral Diencephalon) were included in the analysis (see *Supplementary Figures 1A and 1B*),

### Mice

#### Image Acquisition

Mice where brought under terminal anesthesia using Euthasol 20% (AST Farma) and transcardially perfused with 0,9% NaCL followed by 4% paraformaldehyde (PFA) containing 2 mM PROHANCE (Bracco Imaging Deutschland GmbH). Skulls were collected immediately after perfusion and post-fixed in 5% PFA with 2 mM ProHance for 48 h. Prior to imaging, the skulls were immersed in Fluorinert FC-40 (3M). Ex vivo MRI was performed on a 7-T small-animal scanner (PharmaScan 70/16, Bruker BioSpin GmbH, Ettlingen, Germany) using a 23-mm transmit–receive volume coil.

Imaging was conducted by the Applied Molecular Imaging Erasmus MC (AMIE), at the Leiden University Medical Centre, using a fat-suppressed 3D FLASH sequence (TR = 35 ms, TE = 3 ms, flip angle = 50°). Images were acquired at 70-µm isotropic resolution (matrix size 256 × 210; field of view 17.92 × 14.70 mm) with one excitation (Nex =1).

#### Image processing: Deformation based morphometry

Deformation-based morphometry was performed using the pipeline developed by the CoBrA Lab^13^. An unbiased study-specific template was generated through iterative registration of individual subjects to a developing template. Voxel level differences in local volume were quantified using relative Jacobian determinants derived from the nonlinear deformation fields.

MRI data were preprocessed using the mouse-preprocessing-v8_pv6_brkraw_nifti.sh script, which performs N4 bias-field correction, adaptive non-local means denoising, brain masking via registration to the DSURQE atlas^26^, and field-of-view standardization. Brain extraction was achieved by applying the mask generated during preprocessing. For all analyses, only nonlinear deformations were considered to minimize the influence of global size differences and individual alignment variability.

## Analysis

### Human Cohorts

#### Inclusions and Quality Control

Participants with repeated imaging data and successful cortical reconstruction at both the baseline and follow-up visit were included in the analysis.

In ABCD, only scans that passed the recommended inclusion criteria (imgincl_t1w_include = 1) were considered and we excluded participants with clinically relevant incidental findings (N^excluded^ = 5), and included only one twin at random (N^excluded^ = 1,170). A total of 6,091 individuals (2,793 female) were available for statistical analysis in ABCD.

In Generation R, scans without complete consent form (N^excluded^ = 3), scans with incidental findings (N^excluded^ = 32), and scans with low image quality ratings (N^excluded^ = 1,038) were excluded. For participants with braces, a dedicated braces scan sequence was acquired; scans for which this sequence was unavailable were excluded (N^excluded^ = 790). Additionally, in the case of twins, we included one twin at random (N^excluded^ = 172). A total of 1118 individuals (592 female) were available for statistical analysis in the Generation R cohort.

#### Covariates and Imputation

To best account for potential confounders of the cerebellar-cortical associations and brain development in general, we included covariates capturing key biological and environmental influences on brain maturation, modeled separately for each cohort. Confounders were measured either at baseline or earlier.

In ABCD, the following variables were included as cofounders: ABCD site, race/ethnicity, parental education, gestational age at birth, maternal high blood pressure during pregnancy (dichotomous), maternal smoking/alcohol/anti-depressant/drug use during pregnancy (dichotomous), sum scores of internalizing and externalizing problems based on the Child Behavior Checklist (CBCL)^27^, and total intracranial volume (eTIV).

In Generation R, the following variables were included as cofounders: parental national origin, maternal education, gestational age at birth, maternal pre-pregnancy BMI, maternal systolic (low: < 120; mid: 120-140: high: >140) and diastolic (low: < 80; mid: 80-90: high: >90) blood pressure during pregnancy, maternal infections during the third trimester (dichotomous), maternal smoking/alcohol/benzodiazepine during pregnancy (dichotomous), sum score of prenatal early life stress (https://github.com/SereDef/cumulative-ELS-score), sum scores of internalizing and externalizing problems based on the Child Behavior Checklist (CBCL)^27^, and total intracranial volume (eTIV). Missing covariate data were imputed using *expectation-maximization* (EM) implemented in the Amelia package in R. Imputation was performed separately for each cohort and timepoint. Nominal variables and ordinal variables were specified and a single imputed dataset was generated per timepoint (m = 1), with bounded imputation applied to stressful life events scores (range: 0–3) in Generation R.

### Statistical Analyses

Statistical analyses were performed using R (version 4.4.1) and MATLAB (version R2024a).

#### Primary Analysis

To examine whether early cerebellar morphology predicts subsequent cortical development, we estimated a series of linear regression models in each cohort separately. For each combination of cerebellar ROIs at baseline (N = 30; 10 ROIs x 3 measures [see Regions-of-interest (ROIs) section]) and cortical ROI (N = 44; cortical and subcortical volumes as well as cortical thickness [see Regions-of-interest (ROIs) section]), we modeled the annualized rate of change in cortical ROIs as the outcome, yielding 1,320 (30 cerebellar ROIs x 44 cerebral ROIs) models per cohort. Each model included biological sex assigned at birth, age at baseline MRI, and baseline cortical value of the respective outcome ROI, as well as all aforementioned covariates for each cohort (see Covariates and Imputation). Multicollinearity among covariates was assessed using *variance inflation factors* (VIF), with all values falling within acceptable limits (VIF < 2). The exception was estimated total intracranial volume (eTIV), which showed multicollinearity with other brain measures.

All continuous predictors, outcomes, and covariates were standardized to allow comparison of effect sizes across regions. To minimize the influence of extreme outliers on brain morphology measures, participants were excluded if any cerebellar or cortical region of interest deviated by more than 5 standard deviations from the cohort mean. This procedure was applied separately for each cohort prior to analysis (Exclusions ABCD N = 241; Generation R N = 34). Furthermore, analyses were restricted to right-handed participants to minimize potential confounds of cerebral lateralization. Left-handed participants (ABCD N = 540; Generation R N = 116) were excluded, due to insufficient sample size for model convergence for a left-handed participants only analysis.

Statistical significance was assessed using *false discovery rate* (FDR-BH) correction applied across all 1,320 models within each cohort. An association was considered significant only if it survived FDR correction in both cohorts independently. All analyses were run twice: once with eTIV included in the model as the primary analysis (main text), and once without eTIV included as an additional covariate because of potential effects of multicollinearity, with the latter reported in the supplementary materials.

The final analysis sample for the ABCD cohort included 5,310 participants (2,472 female) and for the Generation R cohort 968 (521 female).

#### Directionality Analysis

To assess the directionality of the cerebellar-cortical relationship, we repeated the full primary analysis in reverse, predicting the annualized rate of change in each cerebellar ROI (n = 30) from each cortical ROI at baseline (n = 44), yielding 1,320 models per cohort. All model specifications, covariates, standardization procedures, and FDR correction thresholds were identical to the primary analysis.

### Mice

#### Statistical Analysis

Voxel-wise linear regression models were applied with sex and group (lesion, sham, and non-surgical) as covariates using the RMINC package in R. *False Discovery Rate* (FDR) was used for multiple testing correction and significant voxels are displayed as t-values at both the 1st and 5th percentile thresholds. To make the identification of structures easier, contours are projected over the unbiased template through the registration of the DSURQE atlas^26^ to this template (Supplementary Figure 2). These regions are also used when reporting the ratio of significance voxels in a given structure, with relabeled regions to match the more commonly used Allen brain atlas.

