## Supplementary Materials for "Cerebellar influences on neocortical development in humans and mice"

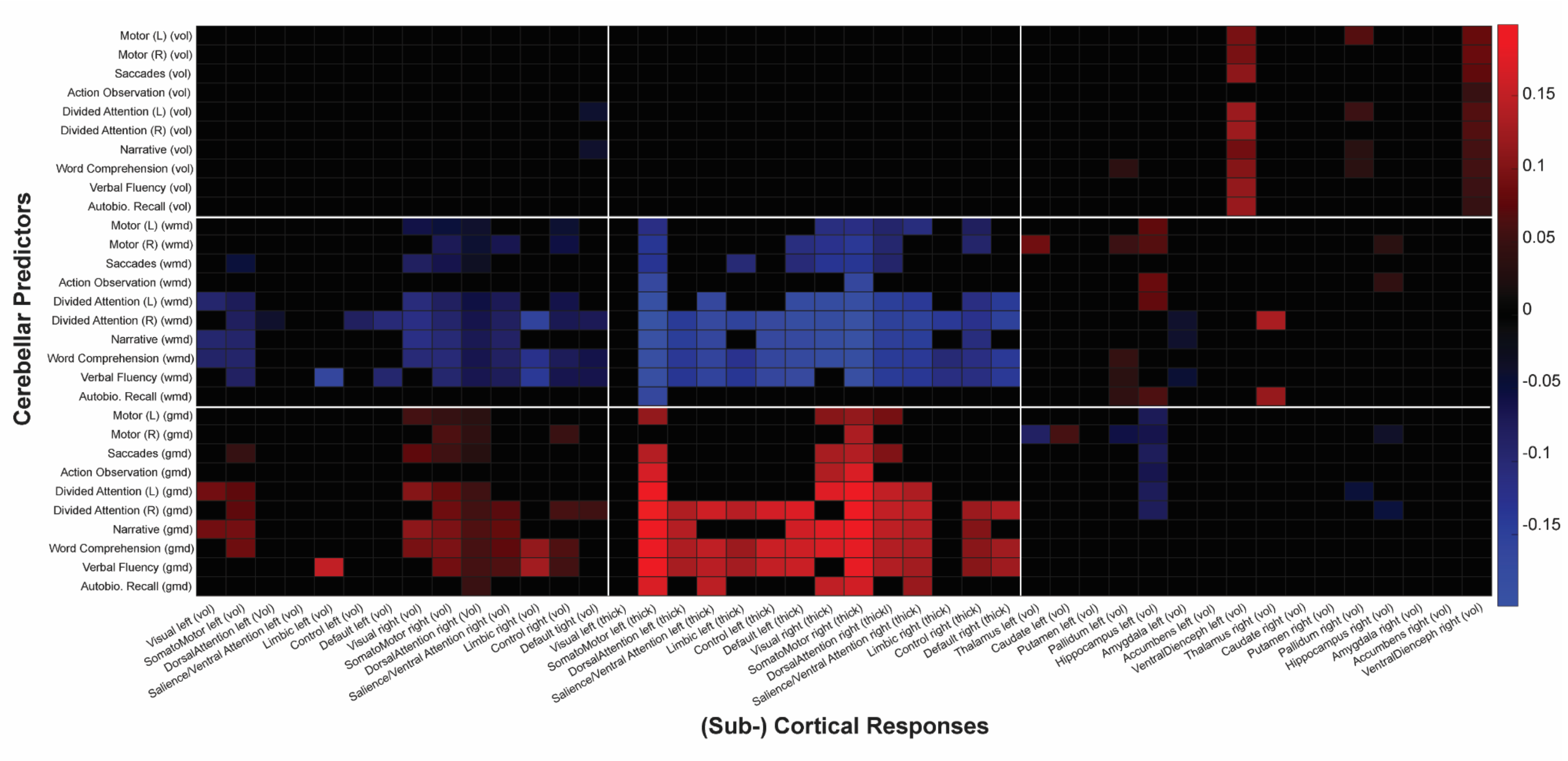


**Supplementary Figure 1A.** Overview of beta coefficients of all models corrected for total brain volume (eTiv)

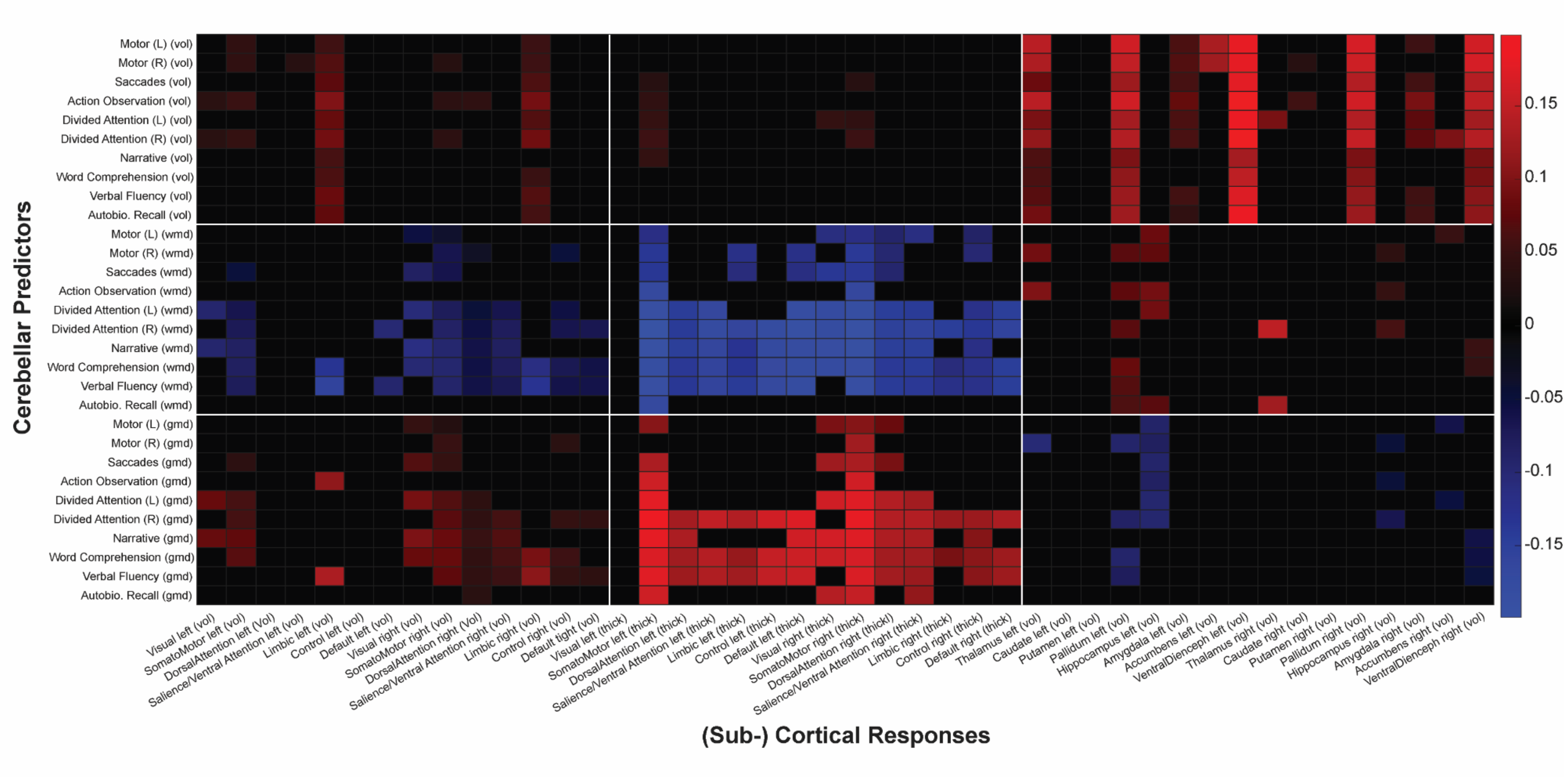


**Supplementary Figure 1B**. Overview of beta coefficients of all models without total brain volume (eTiv) correction

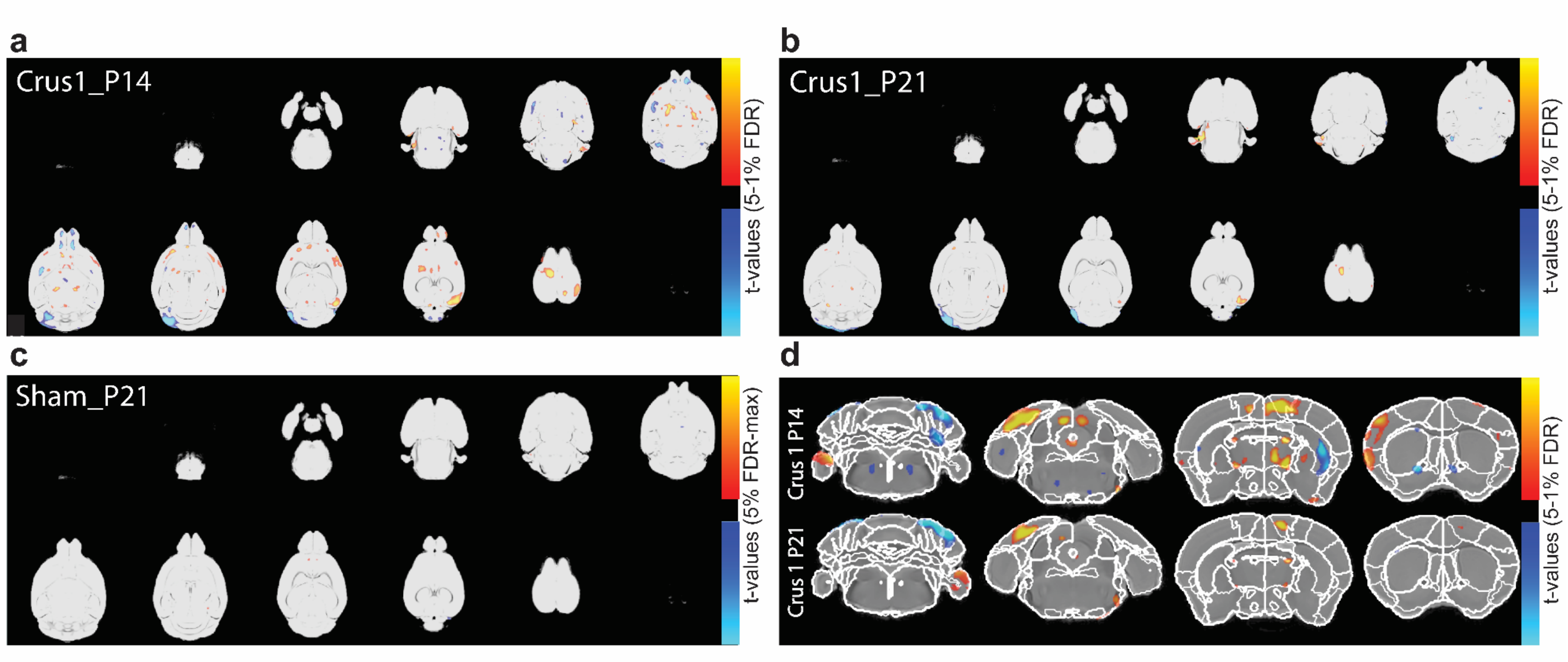


**Supplementary Figure 2.** Significant voxels (after FDR correction) in axial slices for **a.** Crus1 P14 lesion, **b.** Crus 1 P21 lesion, and **c.** Sham P21 (Note: Sham P 14 group is not shown since no voxels survived significance testing). **d.** Atlas overlays in coronal slices for P14 and P21 lesion groups

**Supplementary Table 1.** FDR corrected thresholds for group and sex.

| FDR percentile | F-statistic | tvalue-(Intercept) | tvalue-Groupcrus1_p14 | tvalue-Groupcrus1_p21 | tvalue-Groupsham_p14 | tvalue-Groupsham_p21 | tvalue-Sex |
|---|---|---|---|---|---|---|---|
| 0.01 | 8.04 | 3.17 | 4.42 | 4.53 | NA | NA | 4.65 |
| 0.05 | 4.87 | 2.43 | 3.22 | 3.59 | NA | 4.40 | 3.77 |
| 0.1 | 3.87 | 2.06 | 2.70 | 3.12 | 3.75 | 3.64 | 3.29 |
| 0.15 | 3.31 | 1.83 | 2.37 | 2.83 | 3.25 | 3.23 | 2.95 |
| 0.2 | 2.92 | 1.64 | 2.12 | 2.58 | 2.94 | 2.92 | 2.70 |

**Supplementary Table 2.** Crus 1 Lesion P14 voxel information for each significant structure sorted by highest ratio of significant voxels

| Structure name | Pos sig voxels | Neg sig voxels | Total structure voxels | ratio significant voxels |
|---|---|---|---|---|
| right simple lobule white matter | 0 | 232 | 240 | 0.966666667 |
| right anterior lobule white matter | 0 | 390 | 582 | 0.670103093 |
| left Temporal association areas | 1843 | 0 | 3150 | 0.585079365 |
| right Simple lobule | 0 | 3059 | 5408 | 0.565643491 |
| Nodulus (X) | 0 | 3098 | 5834 | 0.531025026 |
| left Supplemental somatosensory area | 1554 | 26 | 3562 | 0.443571028 |
| right Flocculus | 0 | 245 | 572 | 0.428321678 |
| right crus 1 white matter | 0 | 183 | 429 | 0.426573427 |
| right Cingulate cortex: area 24a' | 334 | 0 | 840 | 0.397619048 |
| left Anterior olfactory nucleus | 0 | 1398 | 3546 | 0.394247039 |
| left Primary somatosensory area, upper limb | 321 | 0 | 872 | 0.368119266 |
| fourth ventricle | 0 | 23 | 66 | 0.348484848 |
| Pyramus (VIII) | 0 | 2085 | 6389 | 0.326342151 |
| lobule 10 white matter | 0 | 172 | 537 | 0.320297952 |
| left Lateral visual area | 1201 | 0 | 3763 | 0.319160244 |
| right Cingulate cortex: area 24b | 194 | 0 | 692 | 0.280346821 |
| right crus 2 white matter | 0 | 141 | 538 | 0.262081784 |
| left trunk of crus 2 and paramedian white matter | 73 | 31 | 409 | 0.254278729 |
| left Primary visual area | 809 | 0 | 3375 | 0.239703704 |
| right Crus 1 | 0 | 1313 | 5813 | 0.225873043 |
| left Secondary visual cortex: mediomedial area | 1345 | 0 | 5990 | 0.224540902 |
| right Cingulate cortex: area 24a | 388 | 0 | 2128 | 0.182330827 |
| right anterior commissure, temporal limb | 169 | 0 | 947 | 0.178458289 |
| left Primary somatosensory area-other | 88 | 0 | 495 | 0.177777778 |
| right Secondary motor area | 866 | 0 | 4987 | 0.173651494 |
| right Primary somatosensory cortex: trunk region | 528 | 0 | 3052 | 0.173001311 |
| right fasciculus retroflexus | 186 | 0 | 1080 | 0.172222222 |
| right PoDG | 203 | 0 | 1244 | 0.16318328 |
| right Insular region: not subdivided | 1667 | 0 | 10350 | 0.161062802 |
| left Primary somatosensory area, lower limb | 1129 | 0 | 7059 | 0.159937668 |
| left Interposed nucleus | 0 | 236 | 1565 | 0.150798722 |
| right Frontal pole, cerebral cortex | 364 | 0 | 2459 | 0.148027654 |
| left Lateral parietal association cortex | 23 | 0 | 170 | 0.135294118 |

| | | | | |
|---|---|---|---|---|
| right Frontal cortex: area 3 | 1116 | 0 | 8377 | 0.133221917 |
| left Medial parietal association cortex | 392 | 0 | 2997 | 0.130797464 |
| right Olfactory tubercle | 273 | 3410 | 30041 | 0.122599115 |
| right Superior colliculus, sensory related | 0 | 48 | 398 | 0.120603015 |
| left middle cerebellar peduncle | 0 | 1529 | 12838 | 0.119099548 |
| left Olfactory areas-other | 1 | 405 | 3492 | 0.11626575 |
| left cerebal peduncle | 438 | 0 | 3776 | 0.115995763 |
| left Agranular insular area, dorsal part | 1051 | 0 | 9159 | 0.114750519 |
| left fasciculus retroflexus | 108 | 0 | 1061 | 0.101790763 |
| left Copula pyramidis | 393 | 60 | 4656 | 0.097293814 |
| left paraflocculus white matter | 308 | 0 | 3226 | 0.095474272 |
| left Primary auditory area | 192 | 0 | 2165 | 0.088683603 |
| right internal capsule | 98 | 0 | 1124 | 0.087188612 |
| right Pallidum, ventral region | 394 | 1 | 4592 | 0.086019164 |
| left Frontal pole, cerebral cortex | 192 | 39 | 2688 | 0.0859375 |
| left Fastigial nucleus | 2 | 198 | 2334 | 0.085689803 |
| left Main olfactory bulb, mitral layer | 45 | 95 | 1845 | 0.075880759 |
| right Paraflocculus | 0 | 59 | 802 | 0.073566085 |
| left subiculum | 312 | 73 | 5517 | 0.069784303 |
| right trunk of crus 2 and paramedian white matter | 0 | 34 | 493 | 0.068965517 |
| right cerebal peduncle | 279 | 0 | 4050 | 0.068888889 |
| right Mammillary body | 1694 | 64 | 25574 | 0.068741691 |
| lobule 1-2 white matter | 0 | 134 | 1994 | 0.067201605 |
| right Anteromedial visual area | 66 | 65 | 1982 | 0.066094854 |
| right medial lemniscus | 0 | 54 | 822 | 0.065693431 |
| right Anterior olfactory nucleus | 0 | 225 | 3533 | 0.063685253 |
| right Cortex-amygdala transition zones | 73 | 0 | 1162 | 0.062822719 |
| right Entorhinal area, medial part, ventral zone | 70 | 0 | 1115 | 0.062780269 |
| right Main olfactory bulb, granule layer | 0 | 440 | 7024 | 0.062642369 |
| right Main olfactory bulb, outer plexiform layer | 0 | 543 | 9180 | 0.059150327 |
| left Thalamus | 0 | 727 | 12421 | 0.058529909 |
| right Secondary visual cortex: mediomedial area | 372 | 0 | 6485 | 0.057363146 |
| right Thalamus | 122 | 0 | 2162 | 0.056429232 |
| left Main olfactory bulb, outer plexiform layer | 155 | 356 | 9104 | 0.056129174 |
| right GrDG | 159 | 0 | 2840 | 0.055985915 |
| right Main olfactory bulb, mitral layer | 0 | 102 | 1843 | 0.055344547 |
| left Frontal cortex: area 3 | 495 | 0 | 9050 | 0.054696133 |
| left Anteromedial visual area | 85 | 19 | 1913 | 0.054364872 |
| right anterior commissure, olfactory limb | 0 | 22 | 409 | 0.053789731 |

| | | | | |
|---|---|---|---|---|
| left Mammillary body | 1064 | 285 | 25622 | 0.052650066 |
| right Ectorhinal cortex | 61 | 0 | 1163 | 0.052450559 |
| right Primary somatosensory cortex: shoulder region | 260 | 0 | 5011 | 0.051885851 |
| left optic tract | 0 | 38 | 733 | 0.051841746 |
| right Inferior colliculus | 2121 | 0 | 41367 | 0.051272754 |
| left Main olfactory bulb, inner plexiform layer | 24 | 56 | 1597 | 0.050093926 |
| CA3Py Inner | 80 | 0 | 1619 | 0.049413218 |
| left Field CA3, stratum oriens | 49 | 0 | 1001 | 0.048951049 |
| left Orbital area, lateral part | 117 | 0 | 2419 | 0.048367094 |
| right Taenia tecta, dorsal part | 53 | 0 | 1100 | 0.048181818 |
| left Main olfactory bulb, glomerular layer | 58 | 226 | 5912 | 0.048037889 |
| left Primary somatosensory area, mouth | 399 | 0 | 8385 | 0.047584973 |
| right Main olfactory bulb, glomerular layer | 0 | 287 | 6225 | 0.046104418 |
| left Lateral septal complex | 11 | 76 | 1894 | 0.04593453 |
| left lateral ventricle-other | 133 | 0 | 3066 | 0.043378995 |
| left Cingulate cortex: area 24a' | 36 | 0 | 853 | 0.042203986 |
| left anterior commissure, temporal limb | 45 | 0 | 1068 | 0.042134831 |
| left Main olfactory bulb, granule layer | 36 | 250 | 6839 | 0.041818979 |
| right Main olfactory bulb, inner plexiform layer | 0 | 62 | 1504 | 0.041223404 |
| right Medial amygdalar nucleus | 536 | 0 | 13269 | 0.040394905 |
| right Primary motor cortex | 412 | 0 | 10395 | 0.03963444 |
| left Medial amygdalar nucleus | 510 | 0 | 12946 | 0.039394408 |
| left Nucleus accumbens | 162 | 0 | 4128 | 0.039244186 |
| left posteromedial visual area | 67 | 0 | 1714 | 0.039089848 |
| left Endopiriform nucleus, ventral part | 0 | 32 | 859 | 0.037252619 |
| right stria medullaris | 0 | 46 | 1242 | 0.037037037 |
| right Dorsal auditory area | 41 | 0 | 1117 | 0.036705461 |
| right Cortical amygdalar area, posterior part, lateral zone | 36 | 0 | 996 | 0.036144578 |
| right mammillothalamic tract | 0 | 61 | 1702 | 0.035840188 |
| right Agranular insular area, dorsal part | 364 | 0 | 10211 | 0.035647831 |
| left Pontine reticular nucleus | 350 | 223 | 16283 | 0.035190076 |
| right Pallidum, dorsal region | 0 | 9 | 262 | 0.034351145 |
| lobule 8 white matter | 4 | 0 | 117 | 0.034188034 |
| left Pallidum, ventral region | 156 | 0 | 4632 | 0.033678756 |
| left Secondary motor area | 106 | 60 | 5025 | 0.033034826 |
| left Retrosplenial area, dorsal part | 69 | 0 | 2115 | 0.032624113 |
| left Medial septal complex | 0 | 206 | 6431 | 0.032032343 |

| | | | | |
|---|---|---|---|---|
| right Entorhinal area, medial part, dorsal zone | 241 | 6 | 8035 | 0.03074051 |
| right Crus 2 | 0 | 118 | 3988 | 0.029588766 |
| left Cingulate cortex: area 29b | 121 | 0 | 4091 | 0.029577121 |
| right Interposed nucleus | 0 | 45 | 1524 | 0.029527559 |
| right Field CA3, stratum radiatum | 187 | 1 | 6597 | 0.028497802 |
| left facial nerve | 73 | 0 | 2576 | 0.028338509 |
| right Cortical subplate-other | 128 | 273 | 14396 | 0.02785496 |
| right habenular commissure | 10 | 0 | 364 | 0.027472527 |
| right corpus callosum | 107 | 0 | 4015 | 0.026650062 |
| CA3Py Outer | 166 | 0 | 6298 | 0.026357574 |
| right Field CA1, stratum lacunosum-moleculare | 0 | 99 | 3776 | 0.02621822 |
| cerebral aqueduct | 0 | 2 | 78 | 0.025641026 |
| right Bed nuclei of the stria terminalis | 49 | 3 | 2062 | 0.025218235 |
| left Insular region: not subdivided | 295 | 0 | 11722 | 0.025166354 |
| posterior commissure | 99 | 0 | 4100 | 0.024146341 |
| Interpeduncular nucleus | 0 | 9 | 380 | 0.023684211 |
| left Primary somatosensory area, nose | 89 | 0 | 3784 | 0.023520085 |
| left Bed nuclei of the stria terminalis | 50 | 0 | 2214 | 0.022583559 |
| right Field CA3, stratum lucidum | 44 | 0 | 2011 | 0.021879662 |
| right subiculum | 120 | 0 | 5510 | 0.021778584 |
| left Ectorhinal cortex | 23 | 0 | 1057 | 0.021759697 |
| right fimbria | 23 | 0 | 1062 | 0.02165725 |
| left Entorhinal area, medial part, dorsal zone | 69 | 106 | 8453 | 0.020702709 |
| left Cortical subplate-other | 8 | 303 | 15548 | 0.020002573 |
| left Primary somatosensory cortex: dysgranular zone | 15 | 0 | 756 | 0.01984127 |
| left Superior colliculus, sensory related | 244 | 0 | 12532 | 0.019470156 |
| right Supplemental somatosensory area | 0 | 72 | 3701 | 0.019454202 |
| Lingula (I) | 0 | 40 | 2061 | 0.019408054 |
| left Piriform-amygdalar area | 9 | 0 | 489 | 0.018404908 |
| left Field CA1, pyramidal layer | 24 | 4 | 1551 | 0.018052869 |
| Central lobule | 0 | 129 | 7241 | 0.017815219 |
| right Olfactory areas-other | 5 | 53 | 3357 | 0.017277331 |
| right Dentate gyrus, molecular layer | 18 | 0 | 1049 | 0.017159199 |
| right Orbital area, lateral part | 38 | 0 | 2233 | 0.017017465 |
| right Pontine reticular nucleus | 199 | 57 | 16070 | 0.015930305 |
| right optic tract | 67 | 0 | 4400 | 0.015227273 |
| right Caudoputamen | 94 | 45 | 9189 | 0.015126782 |
| left Dorsal auditory area | 0 | 18 | 1190 | 0.01512605 |

| | | | | |
|---|---|---|---|---|
| left dorsal fornix | 18 | 0 | 1202 | 0.014975042 |
| left Olfactory tubercle | 282 | 178 | 30975 | 0.014850686 |
| left lateral olfactory tract, general | 47 | 3 | 3440 | 0.014534884 |
| right Cingulate cortex: area 29c | 44 | 0 | 3031 | 0.014516661 |
| right lateral olfactory tract, general | 46 | 0 | 3408 | 0.013497653 |
| right Field CA1, stratum oriens | 32 | 0 | 2505 | 0.012774451 |
| right Paramedian lobule | 17 | 0 | 1384 | 0.012283237 |
| right Piriform cortex | 173 | 0 | 14661 | 0.011800014 |
| left Field CA1, stratum oriens | 17 | 15 | 2805 | 0.0114082 |
| right Cuneate nucleus | 430 | 167 | 52772 | 0.011312817 |
| left Crus 1 | 0 | 63 | 5671 | 0.011109152 |
| right corticospinal tract-other | 158 | 15 | 15769 | 0.010970892 |
| left Cuneate nucleus | 214 | 675 | 81292 | 0.010935885 |
| left Piriform cortex | 149 | 0 | 13709 | 0.010868772 |
| left Primary motor cortex | 99 | 4 | 9712 | 0.010605437 |
| left stria medullaris | 14 | 0 | 1326 | 0.010558069 |
| left pre-para subiculum | 34 | 0 | 3464 | 0.009815242 |
| left corticospinal tract-other | 133 | 27 | 16407 | 0.009751935 |
| right Field CA1, stratum radiatum | 3 | 35 | 4008 | 0.009481038 |
| right Copula pyramidis | 0 | 39 | 4456 | 0.008752244 |
| left Field CA1, stratum radiatum | 29 | 3 | 3699 | 0.008650987 |
| right Field CA1, pyramidal layer | 13 | 0 | 1503 | 0.008649368 |
| Uvula (IX) | 0 | 41 | 5079 | 0.008072455 |
| left habenular commissure | 0 | 3 | 373 | 0.008042895 |
| right Nucleus accumbens | 25 | 7 | 4089 | 0.007825874 |
| left Perirhinal area | 91 | 0 | 12467 | 0.00729927 |
| left Caudoputamen | 63 | 0 | 9756 | 0.006457565 |
| right Primary somatosensory area, lower limb | 55 | 0 | 8552 | 0.006431244 |
| left Cortex-amygdala transition zones | 6 | 0 | 1025 | 0.005853659 |
| right Ventral auditory area | 0 | 22 | 3780 | 0.005820106 |
| left GrDG | 15 | 0 | 2778 | 0.005399568 |
| right Primary somatosensory area, upper limb | 5 | 0 | 997 | 0.005015045 |
| left Ventral auditory area | 17 | 0 | 3414 | 0.004979496 |
| left Cingulate cortex: area 29a | 12 | 0 | 2997 | 0.004004004 |
| right paraflocculus white matter | 13 | 0 | 3269 | 0.003976751 |
| right Parietal cortex: posterior area: rostral part | 0 | 12 | 3273 | 0.003666361 |
| left Field CA1, stratum lacunosum-moleculare | 14 | 0 | 3820 | 0.003664921 |
| left superior cerebelar peduncles | 2 | 0 | 546 | 0.003663004 |
| left Cingulate cortex: area 24a | 8 | 0 | 2201 | 0.003634711 |

| | | | | |
|---|---|---|---|---|
| left Entorhinal area, medial part, ventral zone | 4 | 0 | 1146 | 0.003490401 |
| right Claustrum: ventral part | 0 | 3 | 865 | 0.003468208 |
| right Dorsal nucleus of the endopiriform | 7 | 1 | 2349 | 0.003405705 |
| left Anterolateral visual area | 10 | 18 | 8927 | 0.003136552 |
| left Primary somatosensory cortex: shoulder region | 17 | 0 | 5468 | 0.003108998 |
| right Anterolateral visual area | 8 | 20 | 9193 | 0.003045796 |
| Folium-tuber vermis (VII) | 0 | 17 | 5865 | 0.002898551 |
| right Medial preoptic nucleus | 17 | 0 | 6258 | 0.002716523 |
| right Medial septal complex | 0 | 16 | 6366 | 0.002513352 |
| left Medial preoptic nucleus | 10 | 0 | 4532 | 0.002206531 |
| left Cingulate cortex: area 24b' | 2 | 0 | 1191 | 0.001679261 |
| left Parietal cortex: posterior area: rostral part | 3 | 0 | 3194 | 0.000939261 |
| right Lateral visual area | 0 | 4 | 4466 | 0.000895656 |
| right Cortical amygdalar area, posterior part, medial zone | 1 | 0 | 1132 | 0.000883392 |
| Declive (VI) | 0 | 3 | 3741 | 0.000801925 |
| left Postpiriform transition area | 1 | 0 | 1302 | 0.000768049 |
| right pre-para subiculum | 0 | 2 | 3596 | 0.000556174 |
| left Cingulate cortex: area 29c | 2 | 0 | 3952 | 0.000506073 |
| right Cingulate cortex: area 29b | 1 | 0 | 3904 | 0.000256148 |
| right Lobules IV-V | 0 | 1 | 8166 | 0.000122459 |

**Supplementary Table 3.** Crus1 Lesion P 21 voxel information for each significant structure sorted by highest ratio of significant voxels

| structure_name | pos_sig_ voxels | neg_sig_ voxels | total_structure_ voxels | ratio significant voxels |
|---|---|---|---|---|
| right anterior lobule white matter | 0 | 303 | 582 | 0.520618557 |
| Nodulus (X) | 0 | 2785 | 5834 | 0.477374014 |
| right Simple lobule | 0 | 2276 | 5408 | 0.420857988 |
| right Paramedian lobule | 435 | 0 | 1384 | 0.314306358 |
| left Supplemental somatosensory area | 946 | 23 | 3562 | 0.272038181 |
| right crus 1 white matter | 0 | 113 | 429 | 0.263403263 |
| right copula white matter | 34 | 0 | 131 | 0.259541985 |
| Pyramus (VIII) | 0 | 1470 | 6389 | 0.230082955 |
| right Copula pyramidis | 744 | 112 | 4456 | 0.192100539 |
| right Crus 1 | 0 | 1098 | 5813 | 0.188886977 |
| right fasciculus retroflexus | 140 | 0 | 1080 | 0.12962963 |

| | | | | |
|---|---|---|---|---|
| right anterior commissure, temporal limb | 96 | 0 | 947 | 0.101372756 |
| right trunk of crus 2 and paramedian white matter | 6 | 43 | 493 | 0.099391481 |
| left Lobules IV-V | 0 | 377 | 4779 | 0.078886796 |
| left Primary auditory area | 160 | 0 | 2165 | 0.073903002 |
| right Cingulate cortex: area 24b | 51 | 0 | 692 | 0.073699422 |
| right Frontal cortex: area 3 | 604 | 0 | 8377 | 0.072102185 |
| left Temporal association areas | 215 | 0 | 3150 | 0.068253968 |
| left Primary visual area | 228 | 0 | 3375 | 0.067555556 |
| right Secondary motor area | 325 | 0 | 4987 | 0.065169441 |
| left fasciculus retroflexus | 62 | 0 | 1061 | 0.058435438 |
| left Anteromedial visual area | 101 | 0 | 1913 | 0.052796654 |
| left anterior commissure, temporal limb | 55 | 0 | 1068 | 0.051498127 |
| right Insular region: not subdivided | 506 | 0 | 10350 | 0.048888889 |
| Culmen-other | 0 | 112 | 2721 | 0.041161338 |
| left lateral ventricle-other | 115 | 0 | 3066 | 0.037508154 |
| left Medial parietal association cortex | 106 | 0 | 2997 | 0.035368702 |
| right internal capsule | 38 | 0 | 1124 | 0.033807829 |
| right simple lobule white matter | 0 | 8 | 240 | 0.033333333 |
| right Crus 2 | 0 | 126 | 3988 | 0.031594784 |
| Pons-other | 33 | 0 | 1096 | 0.030109489 |
| right Cingulate cortex: area 24a' | 24 | 0 | 840 | 0.028571429 |
| right Entorhinal area, medial part, dorsal zone | 138 | 83 | 8035 | 0.027504667 |
| left Primary somatosensory area, upper limb | 22 | 0 | 872 | 0.025229358 |
| left Crus 2 | 0 | 80 | 3594 | 0.022259321 |
| right Cuneate nucleus | 1170 | 0 | 52772 | 0.022170848 |
| left Lateral visual area | 82 | 0 | 3763 | 0.021791124 |
| left Thalamus | 0 | 264 | 12421 | 0.021254327 |
| left subiculum | 104 | 0 | 5517 | 0.018850825 |
| right Field CA3, stratum radiatum | 123 | 0 | 6597 | 0.018644839 |
| right Caudoputamen | 140 | 23 | 9189 | 0.017738601 |
| left Nucleus accumbens | 72 | 0 | 4128 | 0.01744186 |
| right anterior commissure, olfactory limb | 7 | 0 | 409 | 0.017114914 |
| right mammillothalamic tract | 20 | 8 | 1702 | 0.016451234 |
| left Crus 1 | 0 | 91 | 5671 | 0.016046553 |
| left pre-para subiculum | 43 | 0 | 3464 | 0.012413395 |
| left Primary somatosensory area, nose | 44 | 0 | 3784 | 0.011627907 |
| right Mammillary body | 275 | 13 | 25574 | 0.011261437 |
| posterior commissure | 46 | 0 | 4100 | 0.011219512 |
| right corpus callosum | 42 | 0 | 4015 | 0.010460772 |

| | | | | |
|---|---|---|---|---|
| Folium-tuber vermis (VII) | 0 | 57 | 5865 | 0.00971867 |
| Lingula (I) | 3 | 17 | 2061 | 0.009704027 |
| left corticospinal tract-other | 80 | 61 | 16407 | 0.008593893 |
| right Parietal cortex: posterior area: rostral part | 0 | 28 | 3273 | 0.008554843 |
| left Retrosplenial area, dorsal part | 18 | 0 | 2115 | 0.008510638 |
| left cerebal peduncle | 32 | 0 | 3776 | 0.008474576 |
| left Pallidum, ventral region | 39 | 0 | 4632 | 0.008419689 |
| right Medial preoptic nucleus | 52 | 0 | 6258 | 0.008309364 |
| right Supplemental somatosensory area | 0 | 30 | 3701 | 0.008105917 |
| left Superior colliculus, sensory related | 100 | 0 | 12532 | 0.007979572 |
| right Inferior colliculus | 312 | 0 | 41367 | 0.007542244 |
| right crus 2 white matter | 0 | 4 | 538 | 0.007434944 |
| left Lateral septal complex | 0 | 14 | 1894 | 0.007391763 |
| right Ventral auditory area | 0 | 27 | 3780 | 0.007142857 |
| left Medial preoptic nucleus | 32 | 0 | 4532 | 0.0070609 |
| left Mammillary body | 6 | 174 | 25622 | 0.007025213 |
| right trunk of simple and crus 1 white matter | 0 | 2 | 291 | 0.006872852 |
| left middle cerebellar peduncle | 0 | 86 | 12838 | 0.006698863 |
| Central lobule | 0 | 48 | 7241 | 0.006628919 |
| left Cuneate nucleus | 450 | 46 | 81292 | 0.006101461 |
| left Lateral parietal association cortex | 1 | 0 | 170 | 0.005882353 |
| left Medial amygdalar nucleus | 67 | 0 | 12946 | 0.005175344 |
| right Taenia tecta, dorsal part | 5 | 0 | 1100 | 0.004545455 |
| left stria medullaris | 6 | 0 | 1326 | 0.004524887 |
| left Interposed nucleus | 7 | 0 | 1565 | 0.004472843 |
| left Agranular insular area, dorsal part | 40 | 0 | 9159 | 0.004367289 |
| right Entorhinal area, medial part, ventral zone | 4 | 0 | 1115 | 0.003587444 |
| right Lateral visual area | 0 | 15 | 4466 | 0.00335871 |
| right optic tract | 12 | 0 | 4400 | 0.002727273 |
| right Interposed nucleus | 4 | 0 | 1524 | 0.002624672 |
| right Agranular insular area, dorsal part | 26 | 0 | 10211 | 0.002546274 |
| left trunk of crus 2 and paramedian white matter | 0 | 1 | 409 | 0.002444988 |
| left Field CA1, stratum oriens | 6 | 0 | 2805 | 0.002139037 |
| left Primary somatosensory area-other | 1 | 0 | 495 | 0.002020202 |
| left Caudoputamen | 19 | 0 | 9756 | 0.001947519 |
| left Field CA1, pyramidal layer | 0 | 3 | 1551 | 0.001934236 |
| right Cingulate cortex: area 24a | 4 | 0 | 2128 | 0.001879699 |
| lobule 10 white matter | 0 | 1 | 537 | 0.001862197 |
| right Thalamus | 4 | 0 | 2162 | 0.001850139 |

| | | | | |
|---|---|---|---|---|
| left Secondary visual cortex: mediomedial area | 10 | 0 | 5990 | 0.001669449 |
| left fimbria | 2 | 0 | 1286 | 0.00155521 |
| left facial nerve | 0 | 4 | 2576 | 0.001552795 |
| left Entorhinal area, medial part, dorsal zone | 13 | 0 | 8453 | 0.001537916 |
| right Field CA3, stratum lucidum | 3 | 0 | 2011 | 0.001491795 |
| left Field CA3, stratum lucidum | 0 | 3 | 2101 | 0.001427891 |
| right GrDG | 4 | 0 | 2840 | 0.001408451 |
| left Bed nuclei of the stria terminalis | 3 | 0 | 2214 | 0.001355014 |
| left Pontine reticular nucleus | 21 | 0 | 16283 | 0.001289689 |
| left Copula pyramidis | 0 | 6 | 4656 | 0.00128866 |
| left Fastigial nucleus | 0 | 3 | 2334 | 0.001285347 |
| right subiculum | 7 | 0 | 5510 | 0.001270417 |
| left posteromedial visual area | 2 | 0 | 1714 | 0.001166861 |
| right lateral olfactory tract, general | 2 | 1 | 3408 | 0.000880282 |
| left Primary somatosensory area, lower limb | 5 | 1 | 7059 | 0.000849979 |
| left Cingulate cortex: area 24b' | 1 | 0 | 1191 | 0.000839631 |
| left dorsal fornix | 1 | 0 | 1202 | 0.000831947 |
| right Frontal pole, cerebral cortex | 2 | 0 | 2459 | 0.000813339 |
| right Primary somatosensory area, nose | 0 | 3 | 4302 | 0.00069735 |
| left Field CA3, stratum radiatum | 0 | 4 | 6764 | 0.000591366 |
| right Anteromedial visual area | 1 | 0 | 1982 | 0.000504541 |
| right Pontine reticular nucleus | 7 | 1 | 16070 | 0.000497822 |
| right Primary somatosensory cortex: trunk region | 1 | 0 | 3052 | 0.000327654 |
| left Dorsolateral entorhinal cortex | 0 | 1 | 3465 | 0.0002886 |
| left Field CA1, stratum radiatum | 0 | 1 | 3699 | 0.000270343 |
| left Perirhinal area | 2 | 1 | 12467 | 0.000240635 |
| left Anterolateral visual area | 2 | 0 | 8927 | 0.000224039 |
| left Olfactory tubercle | 4 | 2 | 30975 | 0.000193705 |
| left Cortical subplate-other | 2 | 1 | 15548 | 0.000192951 |
| left Primary somatosensory cortex: shoulder region | 1 | 0 | 5468 | 0.000182882 |
| right Medial amygdalar nucleus | 1 | 0 | 13269 | 7.53636E-05 |
